# 2D U-NET Deep Learning Architecture for Atherosclerotic Lesion Segmentation and Quantification

**DOI:** 10.64898/2026.10.31.763364

**Authors:** William Giles, Zita Venetianer, Nikita Patel, Monique Bautista Neughebauer, Ashley M. Abramson, Andrew T. Granger, Nicholas J Tustison, Weibin Shi

## Abstract

Mice are commonly used for experimental atherosclerosis research, and a common procedure involves morphometric analysis of atherosclerotic lesions on Oil Red O-stained cross-sections of the aortic root. However, manual measurement of atherosclerotic lesion sizes on numerous histological sections is tedious and time-consuming. Here, we developed a deep learning model based on convolutional neural network to quantify atherosclerotic lesions in cross-sectioned aortic roots of mice. Microscopy images were taken from cross-sectioned aortic roots of genetically diverse F2 mice derived from BALB/cJ and LP/J *Apoe* knockout mice and fed 12 weeks of a Western diet. Aortic cross-sections were stained with Oil Red O and hematoxylin and images captured. Atherosclerotic lesions were manually segmented in 2553 images using Zeiss AxioVision. These images were split into 1,533 for training the U-NET model, 510 for validation, and 510 images for testing. The developed model was evaluated for its performance to measure atherosclerotic lesions by comparing with manual measurement in test images. The model achieved a high degree of accuracy, with an Fl score of 0.82, a Jaccard score of 0.72, a Recall of 0.80, a precision of 0.87, and a correlation coefficient value of 0.98. While manual lesion measurement took approximately 5 minutes per image, the model performed the task in ~3.35 seconds, reducing measurement time by 98.9%. The developed 2D U-NET model provides a new platform for rapid and accurate quantification of atherosclerotic lesion sizes on Oil Red O-stained sections.

## 1. Introduction

Atherosclerosis is the pathological basis of coronary artery disease, ischemic stroke, and peripheral artery disease, which kill more Americans than all types of cancer combined [1]. It is a chronic inflammatory disease of large and medium-sized arteries featured by the buildup of lipid-containing plaques within the artery wall. Growing atherosclerotic plaque and its rupture disrupt blood flow, leading to major adverse cardiovascular events, such as heart attack and stroke [2].

Mice are most commonly used as models for experimental atherosclerosis research. Genetically engineered mice, such as *Apoe* and *Ldlr* knockouts *(Apoe*^−/−^, *Ldlr*^−/−^), develop all phases of atherosclerotic lesions seen in humans [3],[4]. Histology is a common methodology used for in vitro characterization and quantification of atherosclerotic lesions. Atherosclerotic plaque burden of mice is often measured on cross sections of the aortic root and sinus stained with Oil Red O [5]. Manual measurement of lesion sizes on numerous sections is tedious and time-consuming, especially for genetic mapping involving hundreds of mice. Thus, there is a clear need for developing an automated method to speed up measurement of atherosclerotic plaques in mice.

Convolutional neural networks are a type of deep learning models that have the ability to process and analyze images, making them best suited for biomedical image classification, recognition, and segmentation processes as required in the medical field [6]. U-NET is a type of convolutional neural network architecture that is ideal for segmenting 2D biomedical images. It consists of two primary components: the contracting path and the expanding path [7]. The contracting path is down convolutional and consists of a series of repeated convolutional and pooling operations. As images are processed through this path, the feature maps get spatially smaller, increasing the “what” and decreasing the “where”. Next, the feature maps are built up into the original image size in the expanding path, where the up-sample representations at each step are concatenated with the corresponding feature maps in the contraction pathway. We have demonstrated success of using the 2D U-NET supervised model to automatically segment abdominal subcutaneous and visceral fat on CT and magnetic resonance images [8,9]. Here, we sought to use a similar strategy to develop an algorithm for automated quantification of atherosclerotic plaques on Oil Red O-stained sections of mouse aortas.

## 2. Results

### 2.1. Performance of the U-NET model

The resulting model takes in histology images and outputs predicted segmentation masks along with the predicted area of each segmented lesion (Fig. 1). The performance of the model in quantifying atherosclerotic lesions was assessed with 510 test images. Multiple metrics, including the F1 score, Jaccard coefficient, Recall, and Precision, were used to assess its performance (Table 2). The model performed well in predicting plaque areas, achieving a Fl score of 0.824 ± 0.117 (Table 2). The Jaccard coefficient, which measures the intersection of manual and predicted lesion areas on the same images, was 0.715 ± 0.148. The False Negative and Positive errors were low, based on the high Recall value of 0.801 ± 0.143 and the Precision value of 0.866 ± 0.100.

**Figure 1.**
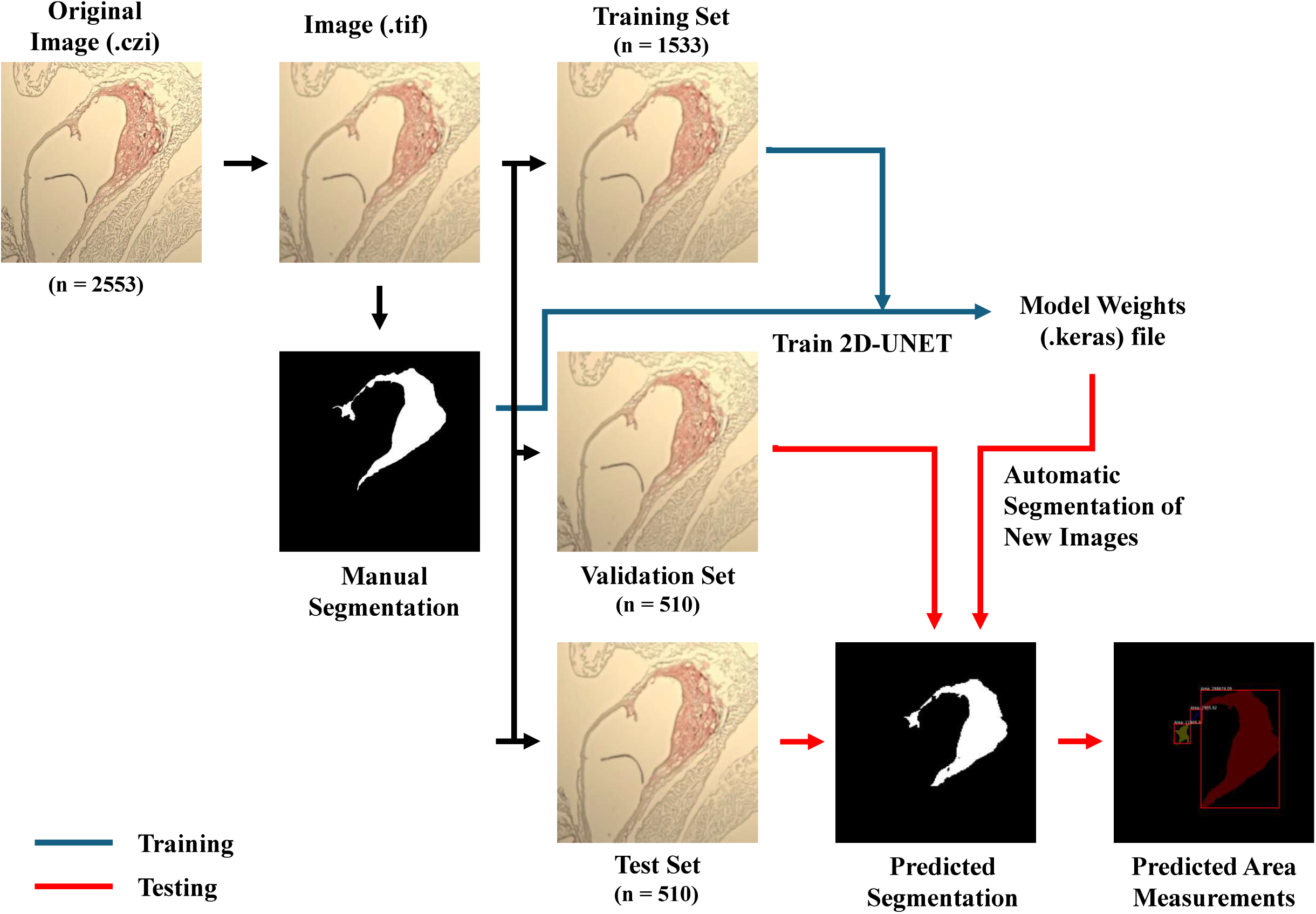
Flowchart illustrating the major steps in developing the deep learning model for quantifying atherosclerotic lesions in aortas of mice. 2553 microscopy images that were manually segmented for atherosclerotic lesions were converted to the .tif format, then resized to 256 × 256 pixels, and finally to binary images. These images were divided into training, validation, and test sets and used for training, validating, and testing the U-NET model.

**Figure 2.**
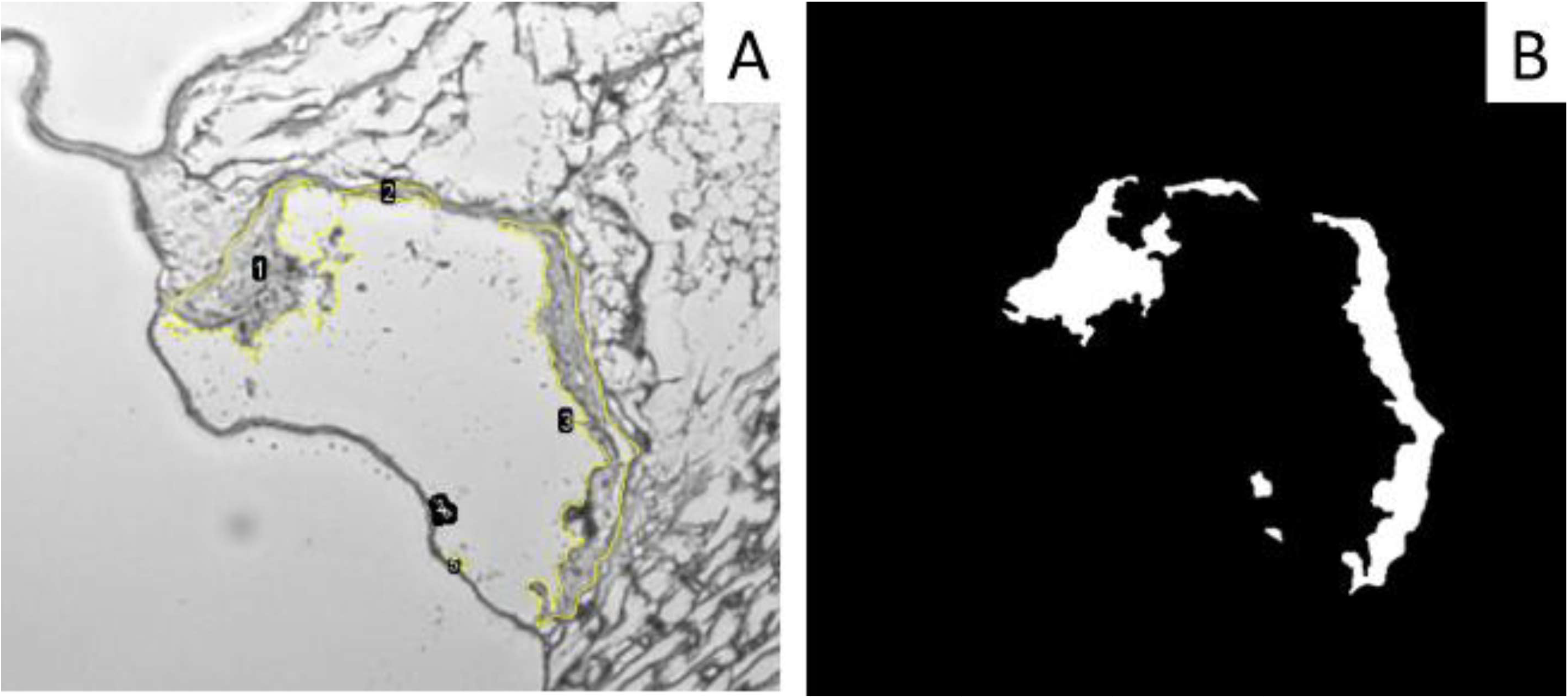
Representative images showing the conversion of a manually segmented image to .tif segmentation masks. The segmentation images in czi format are incompatible with the 2D U-NET architecture. ImageJ was used to transform manual segmentation images in .czi to images in .tif (A) and further to binary masks (B), with white representing the segmented lesions and black everything else. Each Arabic number represents a segmented atherosclerotic lesion, which is encircled by a yellow line. The measurement value for each lesion on the image was saved.

**Figure 3.**
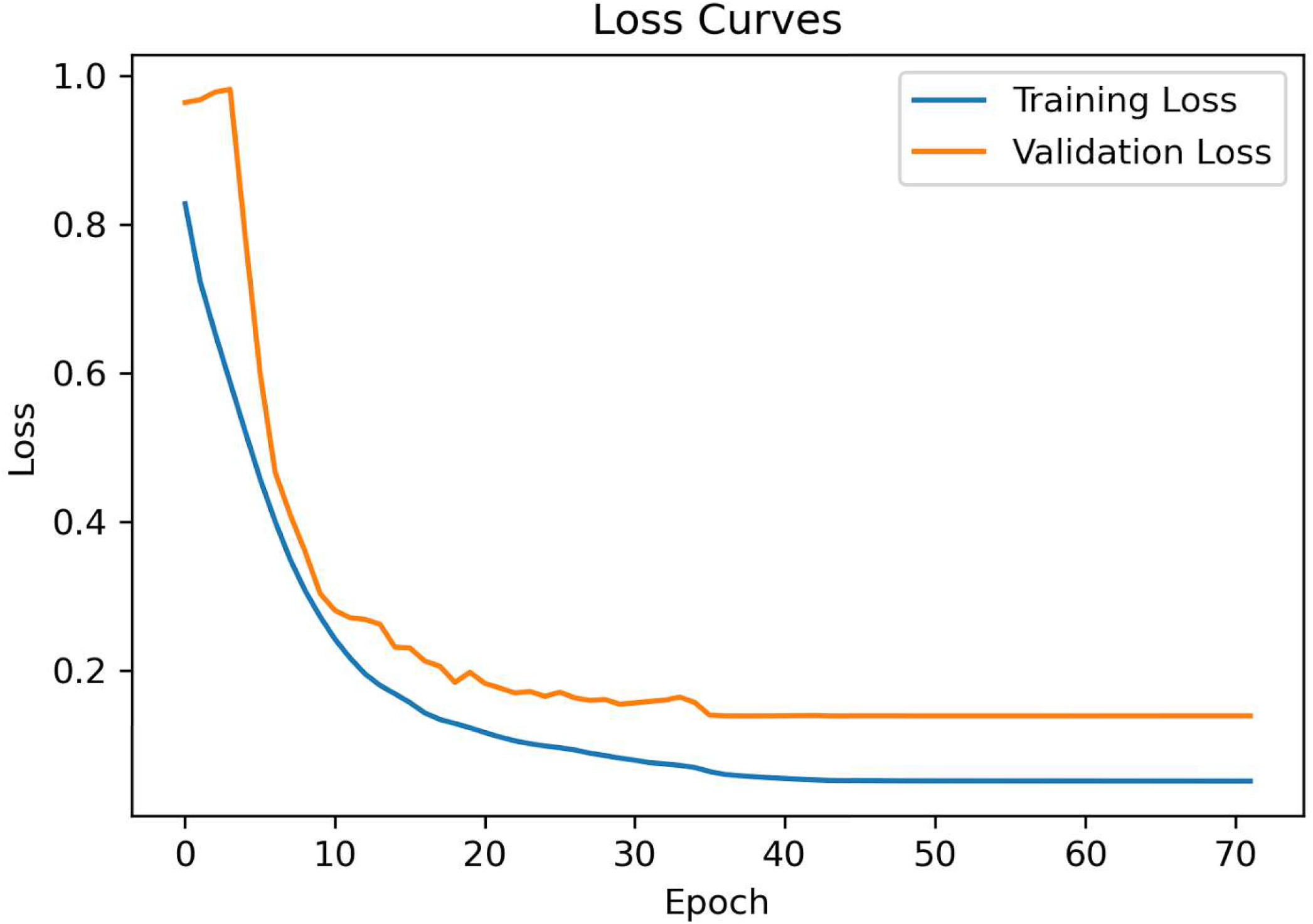
A plot of training versus validation loss during training the U-NET model with the training set. The blue line denotes the training loss (Dice score loss) recorded in successive epochs, and the orange line denotes the validation loss over increasing epochs. Training loss measures how well the model is learning to predict the patterns in the training set. The validation loss measures how well the model can predict on unseen values. A lower loss means that the model is performing better. Both the training loss and the validation loss are stabilized at the 40^th^ epoch. Note no further loss after 40 epochs of training, suggesting that the model has learned the patterns in the training data without overfitting.

The mean lesion area predicted by the U-NET model was 6% smaller than manual measurement for the test images (122,357 ± 91,151 vs. 130,024 ± 91,669 µm^2^) (Fig. 4A). The difference between the two measurements was statistically significant (p = 1.4E-20), suggesting the presence of systemic bias. A Bland-Altman plot helps identify systematic bias and assess the limits of the difference between the two methods. The mean difference is the estimated bias, and the standard deviation (SD) of the differences measures the random fluctuations around this mean [10]. As shown in Fig. 4B, the differences between the two measurements in atherosclerotic lesions are not consistently positive or negative; rather, they distribute on both sides of the solid line representing the mean difference between the two techniques. With few exceptions, the differences between atherosclerotic lesion sizes achieved from the U-NET versus the manual method for the same images are within the mean ± 1.96 SD limits (Fig. 4B), indicating that these differences are still within the normal or standard range and the two methods may be used interchangeably. 4 points on the plot that fall outside of the dash line representing the mean difference− 1.96SD. As outliers have a significant impact on the overall agreement between the two techniques, they were not included in the calculations of regression coefficients and means to prevent them from skewing the results.

**Figure 4.**
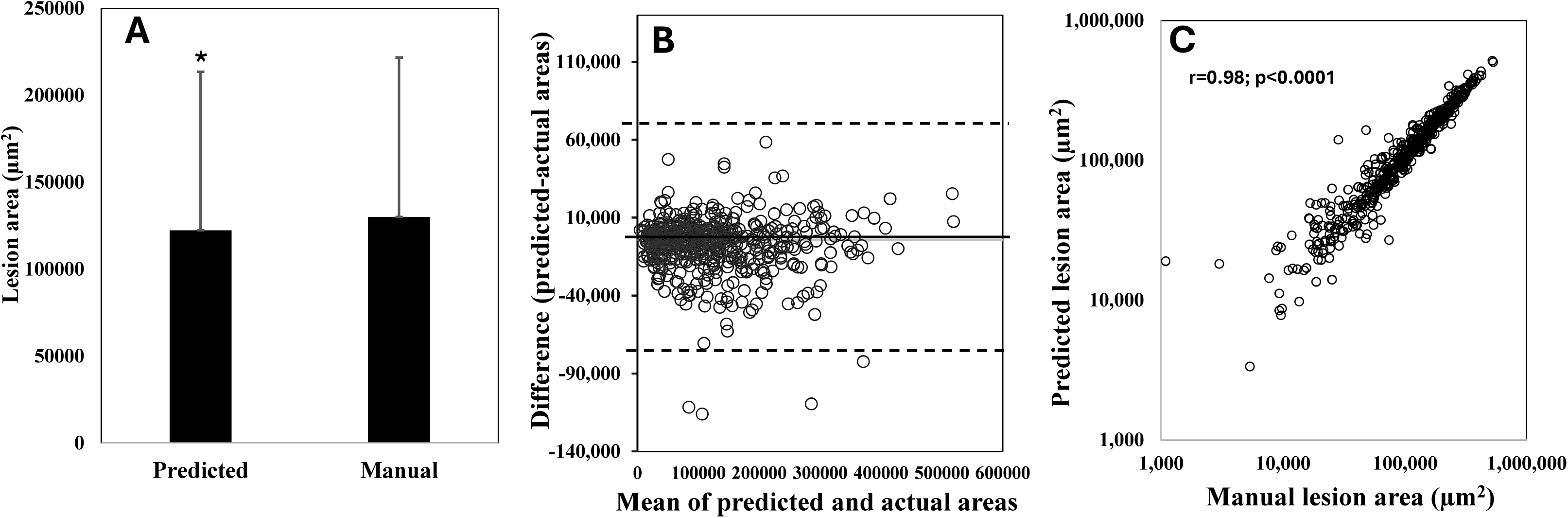
Evaluation of U-NET model performance in segmenting atherosclerotic lesions with the test set. **A**, Average atherosclerotic lesion areas measured by the two methods. Results are means ± SD for 506 test images. 4 outliers with values exceeding the mean ± 1.96 standard deviations (SD) were excluded. **B**, Bland-Altman Plot showing the agreement between two measurement methods by plotting the difference between the two measurements against their average in atherosclerotic lesion sizes for all test images (n = 510). X axis represents the average of the two measurements and Y axis represents the difference between the two measurements on every test image. Each symbol represents a value for a single image. The solid horizontal line denotes the mean difference between the two methods in atherosclerotic lesion areas for all test images. The dash lines denote 1.96 SD above and below the mean difference, respectively. **C**, Correlation between atherosclerotic lesion areas measured by the two methods for the test set. Each symbol represents a value for an individual image. The correlation coefficient (r) and significance (p) are shown.

Regression analysis of associations showed a high agreement in measurements of atherosclerotic lesion areas made by the two methods (r = 0.98; p < 0.0001) (Fig. 4C).

### 2.2. Regional factors affecting model performance

Atherosclerotic lesions vary in shape, size, density, location, and surroundings from mouse to mouse as well as from section to section. Variation in the attributes may impact the performance of the U-NET model. One of the attributes is the lesion size. The model showed a better performance on larger atherosclerotic lesions than smaller ones. As shown in Fig. 5, the percentage error of predicted lesion area compared to manual measurement, calculated as (predicted area − manual area)/manual area) × 100, tends to be smaller with increasing atherosclerotic lesion sizes. Conversely, the percentage error tends to be larger for smaller lesions.

**Figure 5.**
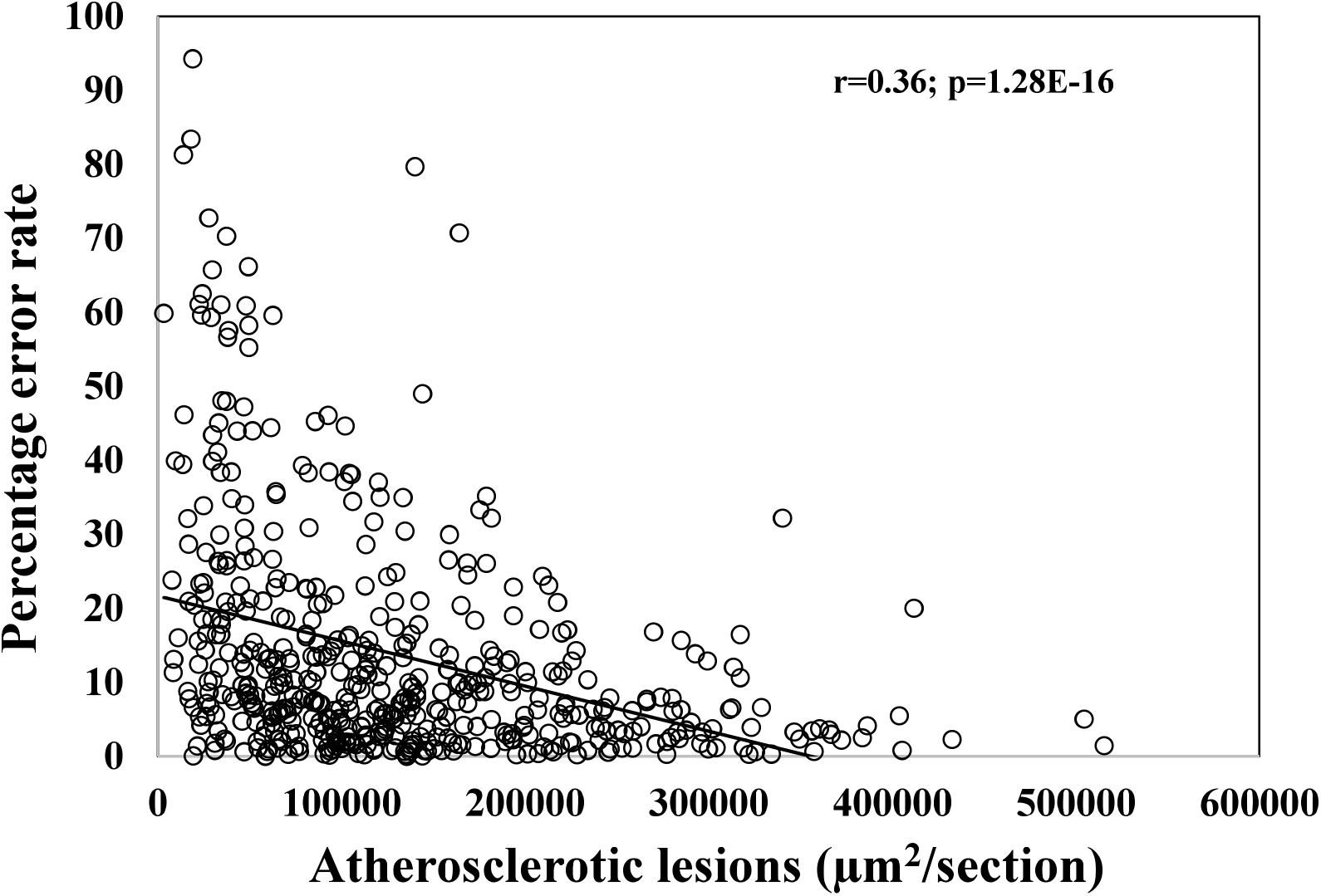
Relationship of the percentage error of predicted to true lesion areas with the true atherosclerotic lesion areas in the test data set (n = 506). Each symbol represents a value for an individual image. The correlation coefficient (r) and significance (p) are shown.

Staining affects the detection of atherosclerotic lesions by the U-NET model. Atherosclerotic lesions are rich in lipids and thus stain red by Oil Red O. However, advanced atherosclerotic lesions, as seen in this study, also contain fibrotic elements, calcification and necrosis, which do not stain red by Oil Red O (Fig. 6). There is fat tissue around the heart and major arteries, known as fat pad, which stains red by Oil Red O. In such cases, the U-NET model may miss non-red stained atherosclerotic lesions (Fig. 6A and C) or count the fat pad as atherosclerosis (Fig. 6B).

**Figure 6.**
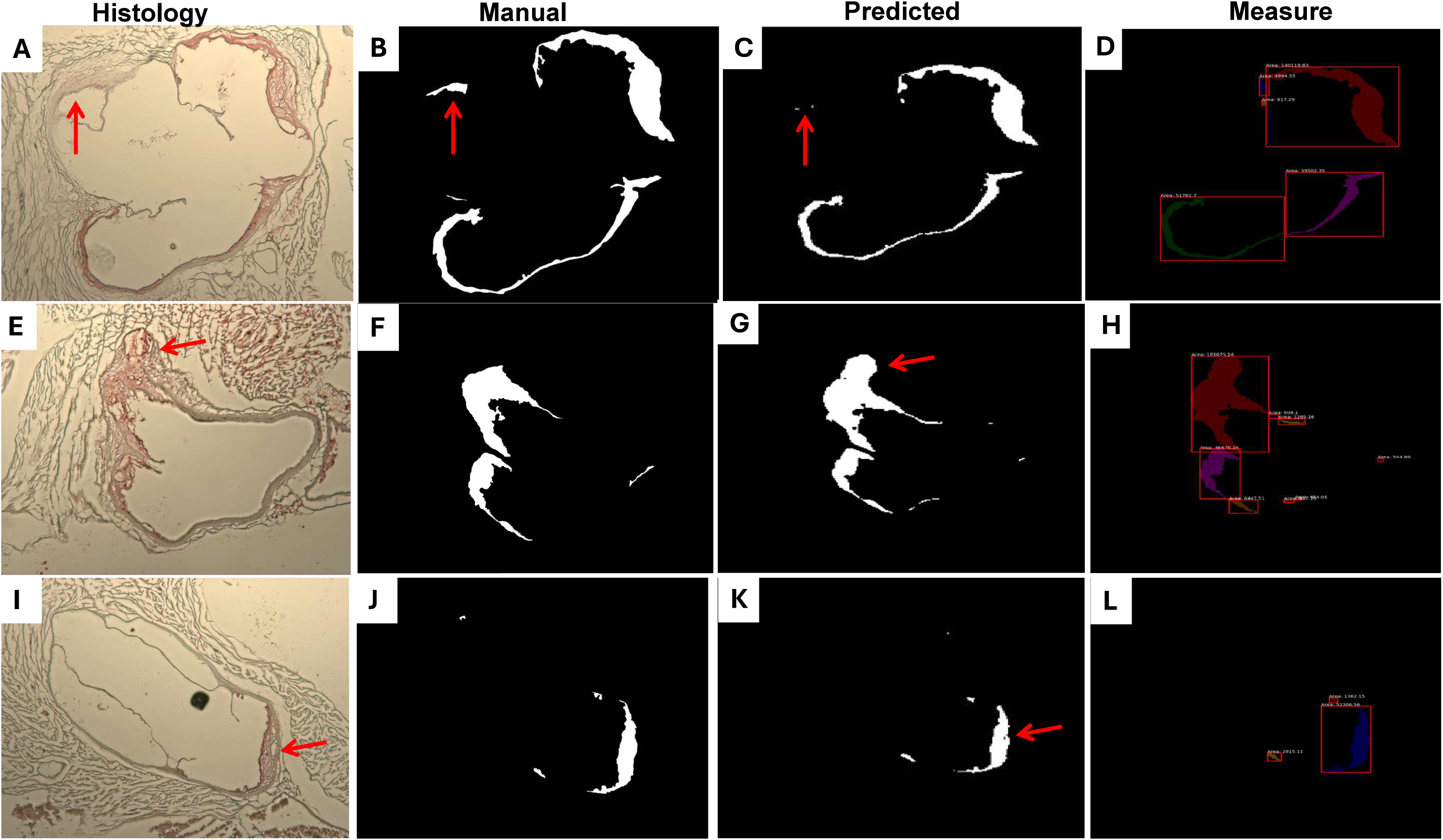
Local factors affecting model performance, including redness of staining (A through D), perivascular fat (E through H), and necrosis (I through L). Red arrows point at affected regions. Under Histology are original microscopy images of aortic cross-sections stained with oil red O; under Manual are binary masks of manually segmented atherosclerotic lesions; under Predicted are binary masks of U-NET segmented atherosclerotic lesions; and under Measure are measurements of U-NET segmented atherosclerotic lesions.

In addition, those lesions with a high prevalence of similar shapes in the training data show high performance scores, probably due to the fact that the model has learned the patterns and relationships that are representative of the real-world reality.

### 2.3. Computational Time

Manual segmentation of atherosclerotic lesion area on one microscopy image took approximately 5 minutes. In contrast, the model inference time was estimated to be 3.350 seconds per image, reducing the time required to segment and measure one image by 98.9%.

## 3. Methods

### 3.1. Histology images

Microscopy images were taken from cross-sectioned aortic roots of F2 mice derived from an intercross between LP/J and BALB/cJ *Apoe*^−/−^ mice [11]. These F2 mice were fed a Western diet for 12 weeks, starting at 6 weeks of age. The procedure for histological processing and staining of aortic root samples was performed as described [12]. Briefly, the aortic root and its adjacent cardiac tissue were harvested, embedded in Tissue-Tek media, and cross-sectioned in 10-µm thickness. Sections were stained with Oil Red O and hematoxylin and counterstained with fast green. Stained sections were photographed under a Zeiss primo star microscope using Zeiss AxioVision software. Captured images were manually segmented to delineate the boundary of each lesion using the AxioVision. Each lesion on segmented 2D images was outlined and had an area measurement in µm^2^. Segmented images were saved as .czi files.

### 3.2. Model development

The overall process of creating an automated segmentation model consisted of data preprocessing, model implementation, model training and tuning, model validation and testing, and model deployment (Fig. 1).

Several image processing steps were taken to ensure the compatibility of manually segmented images with the 2D U-NET architecture. First, segmentation images in .czi were converted to .tif (Fig. 2A) and further to binary masks (Fig. 2B) using ImageJ, with the white area representing the segmented lesion and the black for everywhere else. This step is crucial for training a 2D U-NET, as it ensures the mask to be split in values of 1s and Os, allowing for correct matrix calculations. Additionally, the measurement values on each image were saved.

Then, the images were resized while their original aspect-ratio was maintained. To do this, the cv2 python library was used to reduce the resolutions of images by resizing them and adding padding to ensure that the aspect ratios were not affected. This resizing function was applied to all images in the dataset.

A total of 2553 images was divided into training, validation, and testing sets: 1533 images for training the 2D U-NET, 510 images for validation, and 510 images for testing the model’s performance.

After preparing the histology images and their segmentations, the model implementation process began. The 2D U-NET model was implemented in Python using the Keras library. A custom training script was used to train the model on the prepared datasets.

The 2D U-NET consists of encoder blocks, each containing convolutional layers with batch normalization and ReLU activation, followed by a max-pooling layer for downsampling. At the core of the model is a block that applies convolutional layers with batch normalization and ReLU activation. The model also includes decoder blocks, each consisting of an upsampling layer (transposed convolution), concatenation with the corresponding encoder block’s output, and convolutional layers with batch normalization and ReLU activation. The final output layer is a convolutional layer with a sigmoid activation to produce the segmentation mask. The model is designed to take an input shape of (256, 256, 3) (256 × 256 pixels with 3 color channels) and outputs a binary segmentation mask of the same spatial dimensions.

We did training using the training set and did evaluation on the validation set. Specifically, the model was trained for 100 epochs using a batch size of 32 images and a learning rate of 0.001 using Adam Optimizer.

To select the best epoch model, learning curves were used as a diagnostic tool to monitor the model’s learning based on the training dataset and the validation dataset during training and plotted to show changes in learning performance over time. As shown in figure 3, the training and validation losses decreased with increasing epochs and then reached a point of stability at the 40^th^ epoch. Training stopped after 70 epochs as training and validation losses remained unchanged after 40 epochs. In addition, the gap between the validation loss and the training loss were small at the point of stability, indicating that the model generalized well and was ready for inference.

Throughout training, the validation set served as a benchmark to evaluate whether the model was learning and capable of accurately generalizing prediction from unseen data while avoiding overfitting to just the training images. As shown in Table 1, both training and validation losses were decreasing and the Dice coefficient was increasing over epochs till they reached a plateau at the 40th epoch. The training stopped after 70 epochs as both training and validation losses stopped changing after the 40^th^ epoch of training.

**Table 1.**
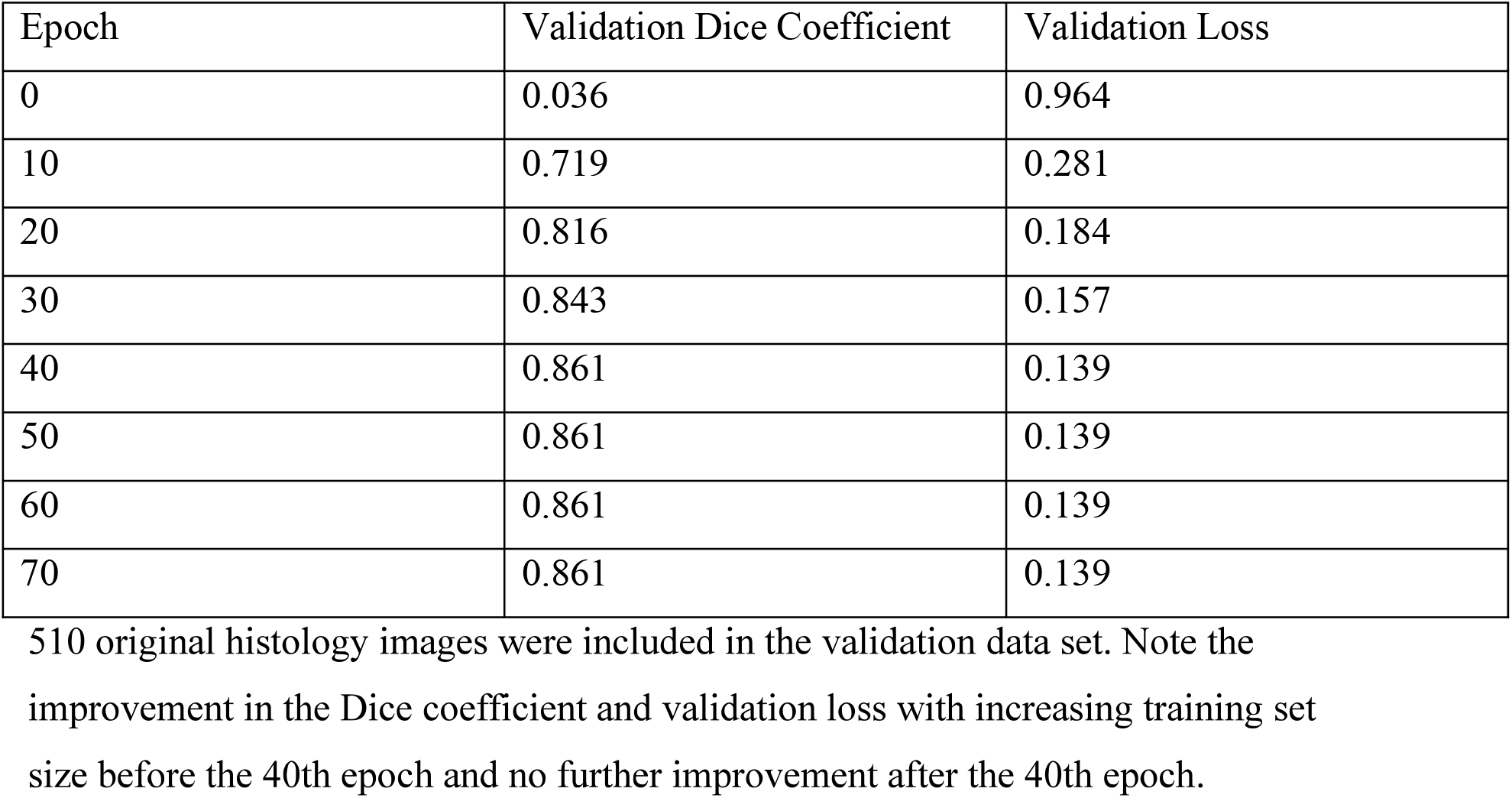
Evaluation of the model’s performance during training on the validation set.

**Table 2.** Performance of the U-NET model in quantifying atherosclerotic lesions on histology images.

|  | F1 Score | Jaccard | Recall | Precision |
|---|---|---|---|---|
| Results | 0.824 ± 0.117 | 0.715 ± 0.148 | 0.801 ± 0.143 | 0.866 ± 0.100 |

The results are means ± SD of 510 testing images.  These measures are used to compare the similarity of the values of atherosclerotic lesion areas measured by the manual and deep learning methods on the same histology images.

After training, the model was evaluated on a separate test set of the original data. It successfully generated binary masks for the test images (Figure 1). After obtaining these binary masks, the scikit-image library was used to calculate measurements for each segmented region in the masks.

The process of obtaining lesion areas from the model-generated binary masks involves identification of separate regions in the binary mask, where there are no overlapping lesion pixels, defined as absence of connected components. Before lesion measurement, the scikit image library is used to fill holes in the binary mask. Then, the scikit image library calculates the area of each lesion, and these calculations are rescaled to match the original manual measurements in µm^2^.

Finally, the model trained can produce binary segmentation masks from raw, histology images, which can then be converted to lesion area measurements in µm^2^. The final model pipeline to produce these measurements is as shown below:

>>> image_path = (“path_to_histology_image”)

>>> segmentation_mask = get_mask(image_path)

>>> measure_rois(image_path, mask)

These three commands will generate a segmentation mask, labeled segmentation mask with lesion measurements, and a file with lesion area measurements.

### 3.3. Analysis metrics

Multiple parameters, including the Fl score, Jaccard index, Recall, and Precision, were used to evaluate the performance of the U-NET model by comparing with manually measured lesion areas on the test set of images. Precision measures how many of the “positive” predictions made by the model are correct:

Precision 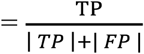, where TP = True Positive; FP = False Positives [13].

Recall or sensitivity measures the model’s ability to correctly identify true positives in the dataset:

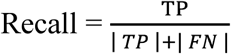, where TP = True Positive; FN = Number of samples wrongly predicted as “negative”.

The Fl score or Dice similarity coefficient combines the precision and recall scores of a model and computes how many times a model has made a correct prediction across the entire dataset:

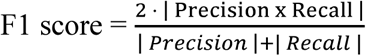. If two segmentations completely overlap, the Fl score is 1; it is 0 if there is no overlap.

The Jaccard coefficient measures the similarity between the predicted data and the true data. For individual labeled region [13]:

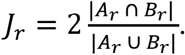

The Jaccard coefficient for all regions:

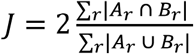

In addition, Pearson’s correlation analysis was performed to determine the correlation between manually measured and predicted lesion areas, as reported [8].

## 4. Discussion

Morphometric analysis of Oil Red O-stained atherosclerotic lesions in the aorta is a common method used to measure atherosclerotic plaque burden of mice. However, manual measurement of atherosclerotic lesions on numerous sections consumes a considerable amount of time. In this study, we developed an automatic segmentation algorithm using the U-NET architecture to quantify Oil Red O-stained atherosclerotic lesions in the aorta of mice. The model has shown notable performance in segmenting atherosclerotic lesions with a Dice score of 0.82, a Recall value of 0.80 and a precision value of 0.87.

Microscopy images were obtained from a genetically diverse F2 population that exhibited a large variation in atherosclerotic lesion sizes, shape and density. Moreover, as atherosclerotic lesions vary in size, shape and density from section to section, images taken from different sections of the same aorta were included in the data set used for model development. Despite the varying attributes of atherosclerotic lesions, the developed model is capable of identifying and segmenting atherosclerotic lesions with a high degree of accuracy. Thus, this model can be applied to the measurement of various atherosclerotic lesions on Oil Red O-stained aortic sections of mice.

The inclusion of a validation data set allowed us to monitor the model’s performance on unseen data during training. We found that both training and validation losses were stabilized at 40 epochs as additional epochs of training led no further decrease in the loss value. This observation suggests that the model was fully trained to a point where further training would not lead to better performance. Thus, the developed model should have learned the underlying patterns and relationships in the training data and can be deployed to measure atherosclerotic lesions on new, unseen data.

The Bland-Altman plot allowed for the visualization of the agreement between the manual and automated methods in segmenting atherosclerotic lesions, any systematic error, and outliers using the test data set. As the differences between the two measurements were not consistently positive or negative, the U-NET model produced no systematic bias in its measurement. With a few exceptions, the differences between the two measurements in atherosclerotic lesions are within the limits of the mean difference± 1.96 standard deviations (SD). This finding suggests that the differences between the two methods of measurements are clinically insignificant and the two methods can be used interchangeably.

Though small, the lesion areas predicted by the U-NET model were smaller than those of manual measurement for the same images. One likely reason is that the new model didn’t count the empty areas within atherosclerotic lesions. As aforementioned, the microscopy images used in this study were obtained from the F2 mice that were fed 12 weeks of Western diet. Thus, these mice developed advanced atherosclerotic lesions [3]. Necrosis and cholesterol clefts are the pathological features of advanced atherosclerotic lesions. Necrotic cellular debris, cholesterol deposits, and even lipids in foam cells can be washed away during staining and form various empty areas within atherosclerotic lesions. As these empty areas are larger than pixels, the U-NET model would count them as non-lesions with pixel values of Os rather than 1s, leading to smaller lesion predictions.

To remedy the flaw, the scikit-image library was implemented to fill empty holes within atherosclerotic lesions. However, when markers aren’t well-defined or boundaries are unclear, scikit-image may miss the regions and leave holes unfilled. Also, this library is sensitive to noise and over-segments when too many minimal holes are present.

We found that many factors affect the performance of the U-NET model, including lesion size, density, component, and perivascular fat. The model showed a higher error rate for smaller atherosclerotic lesions. As the percentage error was calculated as the ratio of the difference between the two measures to the manual measure, smaller lesions will show a larger percentage change than larger lesions for the same amount of change. As discussed above, Oil Red O targets lipids while advanced atherosclerotic plaques also contain non-lipid elements such as necrosis, collagen, and calcification. Perivascular fat stains red with Oil Red O, but it distributes outside the aortic wall. However, the vessel wall can be eroded by atherosclerotic lesions so the separation with the perivascular fat may disappear. In such a case, the model may count the perivascular fat as atherosclerosis.

In summary, we have demonstrated the accuracy of the U-NET model for measuring atherosclerotic lesions on Oil Red O-stained aortic sections of mice. The model can dramatically expedite the process of quantifying atherosclerotic lesions. This model has shown an increased error rate for smaller atherosclerotic lesions. A solution to the problem is to use a higher magnification when small atherosclerotic lesions are photographed.

## Data Availability

The data used for the study can be found in the Supplementary Material file. AtheroQuantNet v1.0 is accessible at https://github.com/wdgstl/Atheroscelerotic_Lesion_UNET.git

## Acknowledgements

Research reported in this paper was supported by NIH grant ROI DK116768. William Giles is a recipient of the Harrison Undergraduate Research Award from the University of Virginia.

